# Respiratory syncytial virus sequesters NF-κB subunit p65 to cytoplasmic inclusion bodies to inhibit innate immune signalling

**DOI:** 10.1101/2020.02.11.944363

**Authors:** Fatoumatta Jobe, Jennifer Simpson, Philippa Hawes, Efrain Guzman, Dalan Bailey

**Affiliations:** The Pirbright Institute, Ash Rd, Guildford, Surrey, UK, GU24 0NF

**Author notes:** Oxford Biomedica (UK) Ltd., Windrush Court, Transport Way, Oxford, UK, OX4 6LT.

## Abstract

Viruses routinely employ strategies to prevent the activation of innate immune signalling in infected cells. RSV is no exception, encoding two accessory proteins (NS1 and NS2) which are well established to block Interferon signalling. However, RSV-encoded mechanisms for inhibiting NF-κB signalling are less well characterised. In this study we identified RSV-mediated antagonism of this pathway, independent of the NS1 and NS2 proteins, and indeed distinct from other known viral mechanisms of NF-κB inhibition. In both human and bovine RSV infected cells we demonstrated that the P65 subunit of NF-κB is rerouted to perinuclear puncta in the cytoplasm, puncta which are synonymous with viral inclusion bodies (IBs), the site for viral RNA replication. Captured P65 was unable to translocate to the nucleus or transactivate a NF-κB reporter following TNF-α stimulation, confirming the immune-antagonistic nature of this sequestration. Subsequently, we used correlative light electron microscopy (CLEM) to colocalise RSV N protein and P65 within bRSV IBs; granular, membraneless regions of cytoplasm with liquid organelle-like properties. Additional characterisation of bRSV IBs indicated that although they are likely formed by liquid-liquid phase separation (LLPS), they have a differential sensitivity to hypotonic shock proportional to their size. Together, these data identify a novel mechanism for viral antagonism of innate immune signalling which relies on sequestration of the NF-κB subunit p65 to a biomolecular condensate – a mechanism conserved across the *Orthopneumovirus* genus and not host-cell specific. More generally they provide additional evidence that RNA virus IBs are important immunomodulatory complexes within infected cells.

**Impact summary:** Many viruses replicate almost entirely in the cytoplasm of infected cells, without too many direct interactions with the nucleus. Examples include respiratory syncytial virus (RSV), measles, Ebola and Nipah; however, how these pathogens are able to compartmentalise their life cycle to provide favourable conditions for replication and to avoid the litany of antiviral detection mechanisms in the cytoplasm remains relatively uncharacterised. In this paper we show that bovine RSV (bRSV), which infects cattle, does this by generating inclusion bodies in the cytoplasm of infected cells. These organelles are unusually membrane-less; likely forming by a process called liquid-liquid phase separation which involves macro-molecular interactions between the viral proteins N and P. We also showed that these organelles, otherwise known as inclusion bodies (IBs), are able to capture important innate immune transcription factors (in this case NF-KB), blocking the normal signalling processes that tell the nucleus the cell is infected. Using fluorescent bioimaging and a combination of confocal and electron microscopy we then characterised this interaction in detail, also confirming that human RSV (hRSV) employs the same mechanism. Like hRSV, bRSV viral RNA replication also takes place in the IB, likely meaning these organelles are a functionally conserved feature of orthopneumoviruses.

## Introduction

Bovine and human respiratory syncytial viruses (bRSV and hRSV, respectively), are closely related viruses that cause acute respiratory illness in cattle and humans, respectively. The viruses infect all ages, but severe illness associated with bronchiolitis and pneumonia is more common in calves (for bRSV) and infants, the elderly and immunocompromised (for hRSV) [1, 2]. Although the process is poorly understood, immune responses to RSV infections are incomplete leading to re-infection, even in healthy adults [3]. In high-risk groups hRSV infection can be fatal; however, there is no approved vaccine and only a single therapeutic option, monoclonal antibodies against the F protein. Whilst there are available bRSV vaccines these are mildly protective and there is evidence for an exacerbation of natural infection [4]. Both viruses were recently taxonomically reclassified as species *Bovine* and *Human orthopneumovirus* within the *Orthopneumoviridae* genus of the *Pneumoviridae* family [5].

bRSV and hRSV are enveloped viruses with a single-stranded negative sense RNA genome, ~15 kb in length, which encodes 11 proteins from 10 mRNAs. Although bRSV and hRSV are restricted to their individual hosts, the viruses and the diseases they cause are similar, making bRSV an excellent model for studying RSV-host interactions. Virus infection and replication within the cell triggers pattern recognition receptors (PRRs) such as toll-like receptors (TLRs) and cytoplasmic nucleic acid receptors (RIG-I and MDA-5), which in turn induce NF-κB- and IRF-dependent signalling [6–8]. NF-κB and IRF are two families of transcription factors that exist as homo- or heterodimers and their activation is regulated at multiple levels. For example, NF-κB p65/p50 dimers are sequestered in the cytoplasm bound to the inhibitor IκBα [9, 10]. Phosphorylation of IκBα by the IκB kinase (IKK) complex targets it for proteasomal degradation releasing p65/p50 for phosphorylation and translocation into the nucleus. Activation and nuclear translocation of IRF-3 homodimers also depends on phosphorylation, through the kinases TBK1/IKKε [11]. Upon activation, these critical transcription factors regulate host cell innate responses, e.g. by inducing cytokines with antiviral activity including type 1 interferons (IFNs), tumour necrosis factor alpha (TNFα) and interleukin-1 (IL-1). Importantly, the mechanisms by which RSV induce or inhibit these signalling pathways is not fully understood.

To overcome this ubiquitous first line of defence, viruses have evolved various inhibitors to modulate these pathways. Viral immune evasion mechanisms include the targeting of receptors, adaptor proteins and/or intracellular kinases in the signalling pathways described above, or indeed directly targeting the transcription factors and their regulators [12, 13], and in this regard RSV is no exception. The RSV SH protein has been shown to be involved in inhibiting NF-κB activation [14, 15], although the exact mechanism of this antagonism is yet to be characterised. As an alternative strategy the RSV NS1 and NS2 proteins have been shown to antagonise IFN-mediated host responses by targeting both type I and III IFN induction [16–18] and signalling [19]. In addition, NS2 interacts with RIG-I inhibiting its interaction with the mitochondrial antiviral-signalling protein (MAVS) [20]. Similarly, NS1 can inhibit phosphorylation of IRF-3 by interacting with MAVS [21]. Recently, the NS proteins have also been shown to be involved in the formation of an “NS-degradasome” that promotes the degradation of components of IFN induction or signalling, including RIG-I, IRF-3, IRF-7, TBK1 and STAT2 [22]. Consequently, activation of the cytotoxic T lymphocyte component of the adaptive immune response is also suppressed [23]. hRSV has also been shown to employ an additional mechanism of innate immune antagonism whereby MAVS and MDA-5 are sequestered into inclusion bodies (IBs), likely through interaction with the RSV nucleoprotein (N protein) [24]. Other cellular proteins involved in the cellular response to viral infection such as, p38 mitogen-activated protein kinase (MAPK) and O-linked *N-acetylglucosamine* transferase (OGT) have also been shown to be recruited into IBs [25].

The cytoplasmic inclusion bodies induced by hRSV infection share many characteristics with liquid organelles or biomolecular condensates [24–26]. They are also structurally and functionally similar to viral inclusions formed by rabies, human metapneumovirus and measles viruses [27–30] and likely represent an essential component of the lifecycle of many negative sense RNA viruses. For the pneumoviruses, these membraneless organelles have been shown to contain N, P, L and M2-1 [26, 29–31]; viral proteins involved in viral genome replication and mRNA transcription, together with the M protein. This presence of viral genomic RNA and mRNA within the IB strongly suggests that these organelles are the primary site for viral RNA replication within the infected cell [26], although this does not appear to be universal a trend, since viral RNA replication of Nipah virus (NiV) was recently shown to occur outside of both of its structurally distinct IB-populations [32]. For RSV and related viruses, ectopic coexpression of the N and P proteins alone results in the formation of IB-like structures, indicating a evolutionarily conserved mechanism for IB formation [24, 26, 28, 30, 31]. Collectively, these data provide strong evidence that events in the bRSV life cycle are not randomly distributed throughout the cell cytoplasm; instead, components of the viral genome, replication machinery and its intermediates are likely to be sequestered away from innate immune sensors in intracellular compartments which are *de facto* viral replication complexes. However, to date there is no evidence on the formation of IBs in bRSV infected cells nor, more broadly, any detailed characterisation of the immunomodulatory effects of the RSV IB on two integral innate immunity transcription factors, NF-κB and IRF3.

Here, we show that in both hRSV and bRSV infected cells, the NF-κB subunit p65 is rapidly sequestered into perinuclear intracytoplasmic puncta. Consequently, activation and nuclear translocation of sequestered NF-κB p65 in response to virus infection and TNFα stimulation are both inhibited. Using both immunofluorescence confocal microscopy and correlative light electron microscopy (CLEM) these puncta were found to be synonymous with the RSV inclusion bodies induced by virus infection. Transmission electron microscopy confirmed that bRSV IBs are not membrane-bound but liquid organelles, likely formed following liquid-liquid phase separation (LLPS). Interestingly, IBs formed by ectopic N and P coexpression were also proficient in colocalising p65. In addition, p65 recruitment was not host-range specific with both human and bovine RSV being capable of sequestering p65, regardless of host cell origin. In addition, we present the first detailed evidence of IB formation in bRSV infected cells, confirming that these viral organelles are the sites of viral RNA replication. Taken together, our data shows an evolutionarily conserved mechanism by which RSV IBs function to compartmentalise viral replication and actively antagonise the innate immune response to infection.

## Results

### BRSV infection induces IRF3, but not NF-κB, nuclear translocation

Given the established role of NF-κB and IRF3 signalling pathways in the cell’s innate response and clearance of viral infection, we used multiple approaches to examine the activation of these transcription factors following bRSV infection. Vero cells were infected with bRSV at a multiplicity of infection (MOI) of 1 for 24h. Cells were then immuno-stained for bRSV F as a marker for infection as well as for the NF-κB subunit p65 or, separately, IRF3. Immunofluorescence (IF) analysis of mock-infected cells confirmed that both transcription factors are normally located in the cytoplasm (Fig 1A). When the NF-κB and IRF3 pathways were stimulated in mock-infected cells with agonist treatment (hTNFα and poly[I:C], respectively), both the NF-κB subunit p65 and IRF3 translocated from the cytoplasm to the nucleus, as expected (Fig 1A top panel; inset zooms). However, although infection with bRSV induced similar levels of IRF3 nuclear translocation (Fig 1A; bottom right panel), significantly the NF-κB subunit p65 remained cytoplasmic, coalescing into intracytoplasmic puncta, mostly perinuclear and present only in infected cells (Fig 1A; bottom left panel). Fluorophore intensity profile analysis was then performed to assess the relative accumulation of both p65 and IRF3 in infected and/or stimulated cells. For IRF3, poly(I:C) stimulation of infected cells enhanced its nuclear translocation, relative to uninfected cells (Fig 1A; bottom right – inset zoom). However, IF and intensity profile analysis revealed that, even in the case of hTNFα stimulation, p65 nuclear translocation in bRSV infected cells was absent and that most p65 remained in the observed perinuclear puncta (Fig 1A; bottom left – inset zoom). bRSV can infect a broad range of host cells *in vitro* – growing to similar titres in both Vero and Madin-Darby bovine kidney (MDBK) cells (supplementary Fig 1A and B). To examine the apparent innate immune antagonism in bovine cells, equivalent infections were performed in MDBK cells. These experiments confirmed the same p65 sequestration into perinuclear puncta following bRSV infection, as well as the related insensitivity to TNFα stimulation (supplementary Fig 1C) indicating a conserved mechanism of antagonism active in both primate and ruminant cells.

**Figure 1.**
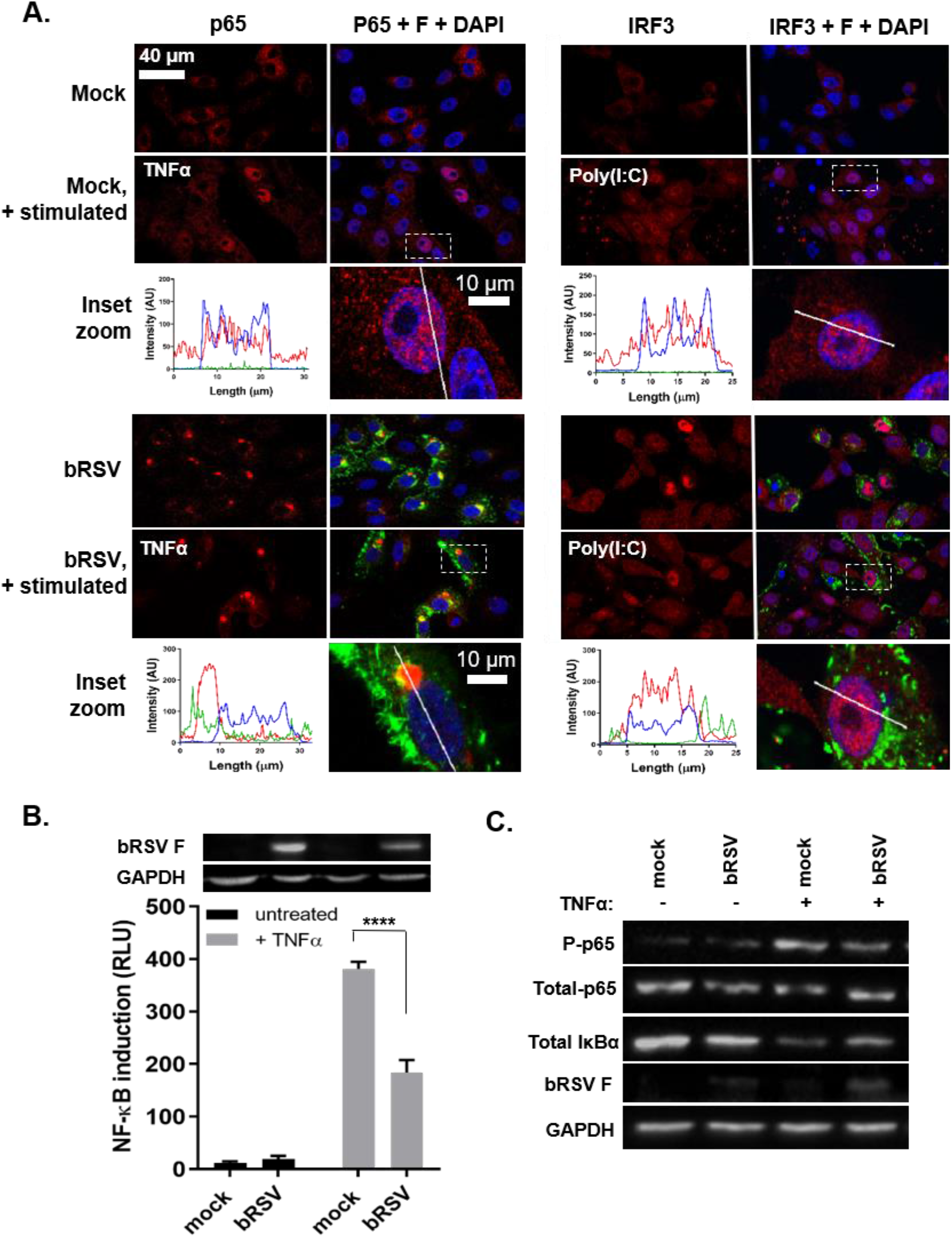
bRSV infection induces IRF3, but not NF-κB, nuclear translocation. **(A)** Vero cells, uninfected (mock), or infected with bRSV at an MOI of 1 for 24 h, were left untreated, stimulated with 20 ng/ml hTNFα for 30 mins or transfected with 2.5 μg/ml poly(I:C) and incubated for 6 hrs at 37°C. Cells were then fixed and immunostained with anti-RSV F (green) and anti-NF-κB p65 or anti-IRF3 (red) antibodies. Cell nuclei were stained with DAPI (blue) and images obtained using a Leica TCS SP5 confocal microscope. The boxed areas are shown magnified in the panels below (inset zoom). Graphs show fluorescent line intensity profiles along the respective white lines within these inset zooms. **(B)** 293T cells were mock infected or infected with bRSV at an MOI of 1. At 6 h p.i., cells were transfected with 100 ng NF-κB FLuc reporter and 10 ng TK-renilla luciferase and incubated at 37°C. At 18 h p.t., cells were left untreated or stimulated for 16 h with 20 ng/ml hTNFα. Cells were then lysed and analysed for firefly and renilla luciferase activities. Graph depicts means ± SD of three replicates from the same experiment. As controls, the levels of RSV F and GAPDH were analysed by western blotting on a fourth replicate. Statistical significance determined by ANOVA as described in the methods, \*\*\*\**p*<0.0001. **(C)** Vero cells mock infected or infected with bRSV at an MOI of 2 for 24 h were left untreated or stimulated with 20 ng/ml hTNFα for 10 mins. Cells were then lysed and analysed by western blotting for phosphorylation of p65 using phospho-specific forms of the antibody, total p65, IκBα and RSV F. GAPDH was detected as a loading control.

To examine the effect of this sequestration on NF-κB signalling, we next employed a luciferase reporter assay to assess NF-κB transactivation. HEK293T cells were infected with bRSV at an MOI of 1, before being transfected with the NF-κB reporter and subsequently treated with or without TNFα (Fig 1B). Interestingly, infection without TNFα treatment did not result in any significant activation of the reporter, despite demonstrable viral protein production (Fig 1B, black bars and RSV F western blot), indicating that even in the presence of active viral replication there is little to no activation of the NF-κB signalling pathway in bRSV-infected cells. Indeed, activation of the NF-κB reporter was only seen following addition of 20 ng/ml of exogenous hTNFα; however, this activation was significantly less in infected cells, when compared to the mock (Fig 1B, grey bars). Separately, we also examined protein levels of p65 (total and transiently phosphorylated) and IκBα, components of NF-κB signal transduction, in infected Vero cells with and without TNFα stimulation. As expected TNFα treatment of mock-infected cells resulted in an increase in p65 phosphorylation and a decrease in total IκBα (presumably the result of proteasomal degradation following its own phosphorylation) (Fig 1C; mock -/+ TNFα) [9]. The detected levels of phospho-NFκB p65 and total IκBα in infected cells (Fig 1C; infected -/+ TNFα) confirmed the lack of activation during infection and also the modest NF-κB activation induced by bRSV infection with subsequent TNFα treatment observed in Fig 1B. Together, the data strongly suggests that NF-κB signalling is inhibited by bRSV infection due to its sequestration into intracytoplasmic puncta. Importantly, these data also indicate that the sequestered p65 is not in a transcriptionally active state, since infection did not result in a marked increase in p65 phosphorylation nor evidence for demonstrable IκBα degradation.

### BRSV replication induces the recruitment of the NF-κB subunit p65 into intra-cytoplasmic bodies distinct from stress granules

NF-κB p65 puncta were only observed in bRSV infected cells showing detectable levels of F protein, indicating a correlation between productive infection and sequestration (Fig 1A). To examine this correlation and define the kinetics of p65 sequestration over time, MDBK cells were infected at an MOI of 1 and fixed at different times post infection (p.i), before being permeabilised and the distribution of p65 and RSV F analysed by IF. Detectable NF-κB p65 puncta (>3 μm^2^) were apparent in infected cells by 16 h p.i. (Fig 2B), correlating with significant levels of F expression (Fig 2A). Interestingly, two populations of F protein were present at this stage, a perinuclear, presumably ER- or vesicle-associated population (Fig 2A; white arrow), and a peripheral more filamentous-like population, possibly the site of virion biogenesis (Fig 2A; beige arrow) – neither of which appeared to colocalise in any significant way with p65. By 24 h p.i., all infected cells contained at least one p65 puncta with none being observed in nearby uninfected cells. Using fluorophore line of interest analysis, we were also able to assess the ratio of cytoplasmic- to puncta-localised p65 as well as the increasing diameter of these aggregates. As infection proceeded the intensity of p65 in the puncta increased as the level of disperse p65 in the cytoplasm decreased (Fig 2C; ‘p65 in puncta’ vs. ‘p65 outside puncta’), indicating coalescence, and supporting our observations in Fig 1C that the total amount of p65 in cells does not dramatically change during infection, only its sub-cellular localisation. Average puncta size increased as infection progressed with p65 aggregations at 48 h p.i. having a mean area of 22.18 μm^2^ (Fig 2B). Smaller p65 puncta (<10 μm^2^) were also observed at 48 h p.i., most likely the result of nascent infections in nearby cells. By this time F protein expression was markedly different, with less distinct populations of protein; however, there was still no obvious co-localisation with the p65 puncta. A similar pattern of results was also observed in Vero cells (supplementary Fig 2).

**Figure 2.**
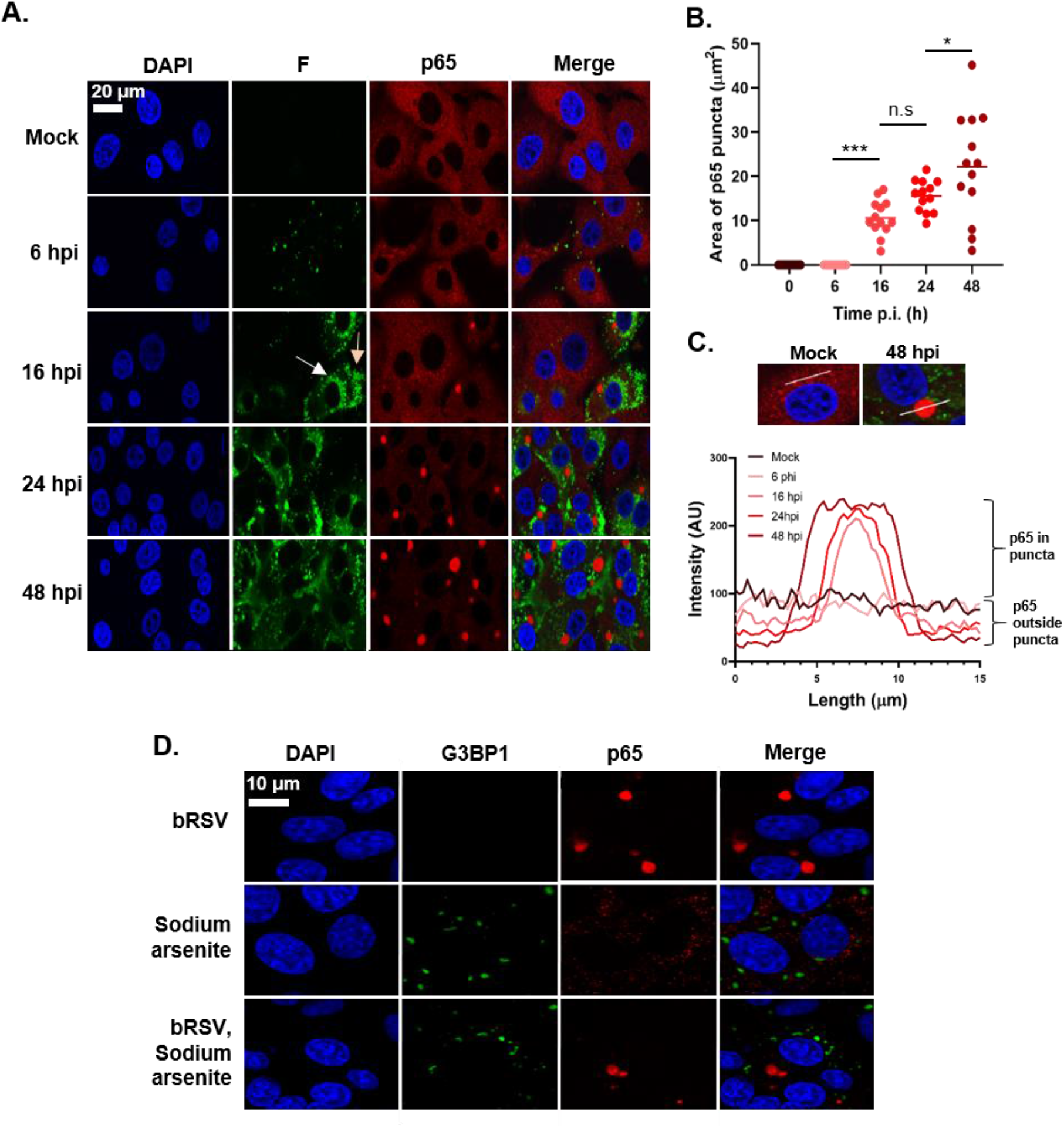
BRSV replication induces the recruitment of the NF-κB subunit p65 into intra-cytoplasmic bodies distinct from stress granules. **(A)** MDBK cells were mock infected or infected with bRSV. At the indicated times p.i. cells were fixed and immuno-stained with anti RSV F (green) and anti-NF-κB p65 (red) antibodies. Nuclei were stained with DAPI (blue) and images obtained using a Leica TCS SP5 confocal microscope. **(B and C)** Quantification of p65 puncta in A obtained using the quantify tool of Leica LAS AF Lite software as described in the methods. **(B)** Surface area of thirteen p65 puncta per time point and mean area are indicated. Statistical significance determined by ANOVA as described in the methods, n.s: non-significant; \**p*<0.05; \*\*\**p*<0.001. **(C)** Graph showing the line intensity profiles along chosen 15 μm lines of interest (example micrographs: 15 μm drawn across a puncta, or, across the cytoplasm in mock cells) of an average of five puncta per time point. **(D)** Vero cells were infected with bRSV or mock infected. At 24 h p.i., cells were treated with 500 μM Sodium arsenite or mock treated for 1 hr. Cells were then fixed and immuno-stained with anti-G3BP1 (green) and anti-NF-κB p65 (red) antibodies. Nuclei were stained with DAPI (blue) and images obtained using a Leica TCS SP5 confocal microscope.

Our first line of inquiry following the identification of p65 puncta in bRSV infected cells was based on their visual similarity to protein and mRNA aggregations that form in cells in response to cellular stress and viral infections, so-called stress granules (SG). A wide range of viruses have been shown to either induce or inhibit SG formation to their advantage [33]; however, there are contradictory findings on SG induction by RSV [25, 34–36]. To examine the potential relationship between these p65 puncta and SG we induced SG formation in bRSV infected cells with sodium arsenite treatment and performed co-immunostaining for p65 and G3BP1 (a SG marker) in fixed cells. Although we were able to successfully stimulate the production of SGs in Vero cells our analysis showed that the p65 puncta were entirely distinct from these granules (Fig 2D). Tangentially, this experiment also demonstrated that bRSV infection does not significantly induce SG formation.

### The NF-κB subunit p65 co-localises with viral inclusion bodies independently of RSV-encoded immunomodulators

RSV has a relatively small genome, encoding 11 proteins from 10 genes (Fig 3A). Recent work has demonstrated that hRSV infection induces the formation of inclusion bodies (IB) which contain components of the RNA polymerase complex and ribonucleoprotein (RNP), notably N and P [26]; however, to our knowledge, similar IBs have not been identified, or functionally characterised, in bRSV-infected cells. To examine the presence of IBs, the distribution of bRSV proteins and, collectively, their sub-cellular localisation in relation to the observed p65 puncta, we infected Vero cells and fixed them, along with mock infected cells, for IF analysis at 24 h p.i. These cells were then co-immunostained for p65 and bRSV N, P, M or F proteins. As expected, neither p65 puncta or bRSV proteins were detected in mock infected cells (Fig 3B). Similarly, as described in Fig 1 and 2 RSV F did not colocalise with p65 or show evidence of sub-cellular localisation with IB-like structures. In contrast, in infected cells, three of the viral proteins (N, P and M) predominately localised to large intracytoplasmic organelles, characteristic of viral inclusion bodies (Fig 3B; green panels), although smaller N-positive IBs were also present (see below). Although there was a varying degree of cytoplasmic signal for N, P and M outside of the IBs, most of the IF signal was found within these structures (Fig 3B; zoomed inset and line of interest plots). The sub-IB localisation of bRSV N and P was similar to that previously described for hRSV, with N and P being found on the periphery of the organelle [26]. The significant intra-IB localisation of the M protein at 24 h p.i., as well as its partial nuclear localisation, is consistent with previously reported IF in RSV-infected cells [37, 38]. However, the role of M in RNA virus IBs reflects an interesting point of divergence; with some viral IBs being M positive (e.g. RSV) and others negative (e.g. rabies) [39]. Significantly, the larger N, P or M-positive IBs were, in the majority of cases, also p65 positive (Fig 3B; red IF panels) identifying, for the first time, that this NF-κB component was being recruited to RSV inclusion bodies in infected cells. To examine this in detail we next characterised the number, size and p65 status of N-positive IBs in infected cells, observing that they were numerous and mostly localised in the median section of the cell. We therefore obtained images from multiple planes in this section to assemble max intensity z-stacks to aid quantification. From 16 h p.i., N and p65 positive IBs were evident throughout the cell in a conserved pattern consisting of a single, large and perinuclear IB with multiple smaller inclusions more evenly distributed through the cytoplasm (Fig 3C, supplementary Fig 3 and supplementary video 1 and 2). Using z-stacks, we quantified the number per cell (counting 18 cells per sample, per timepoint) and surface area of N and/or p65 positive structures >0.1 μm^2^, observing these both increasing as infection progressed. The average number of IBs >0.1 μm^2^ grew from 1.7 per cell at 6 h p.i., to 23.8 at 24 h p.i. (Fig 3D). Their mean area also increased to 8.99 μm^2^ by 24 h p.i. (Fig 3E), significantly influenced by the presence and growth of the larger IB. p65 positive IBs were detected from 16 h p.i.; however, p65 was only detected in larger IBs (>1.39 μm^2^) (Fig 3E) with up to 4 of these being evident per cell (Fig 3D). In conclusion, although multiple N-positive IBs are present in infected cells it is predominantly the larger IBs which contain the sequestered p65. Together, these data suggest that bRSV infection induces the formation of IBs in the cytoplasm of infected cells, organelles which are also involved in sequestering cellular proteins to effect immunomodulation. To our knowledge, this represents an entirely novel mechanism of viral inhibition of NF-κB signalling, since it is the sequestration of signalling components to a viral organelle, rather than the degradation commonly seen [12, 22], which leads to the innate immune antagonism witnessed in Fig 1.

**Figure 3.**
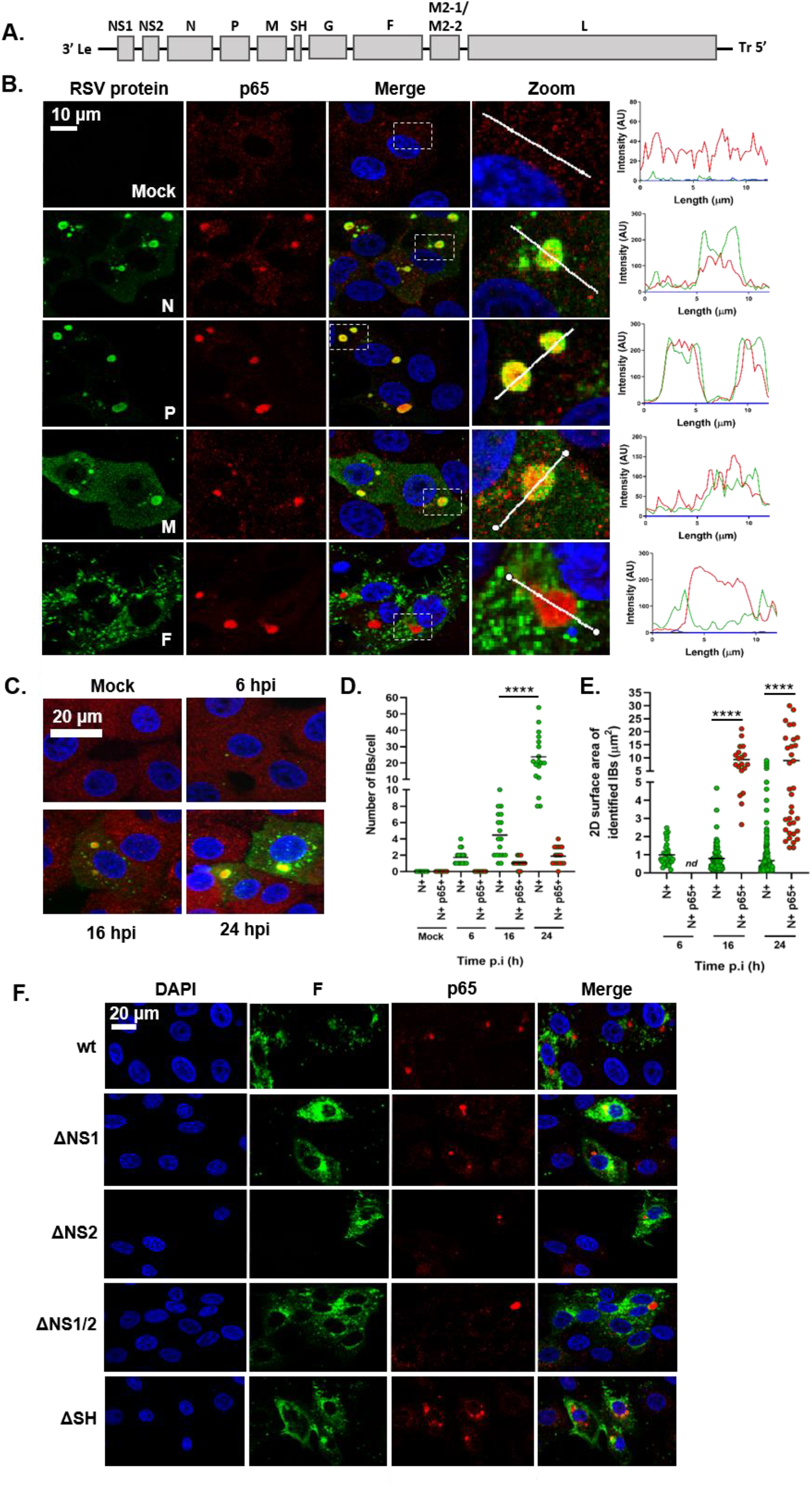
The NF-κB subunit p65 co-localises with viral inclusion bodies independently of RSV-encoded immunomodulators. **(A**) Schematic depiction of the bRSV genome showing organisation of the encoded genes. **(B)** Vero cells, mock infected, or infected with bRSV for 24 h, were fixed and immunostained with rabbit anti-NF-κB p65 (red) and mouse monoclonal anti-RSV N, P, M or F antibodies (green). Nuclei were stained with DAPI (blue) and images obtained using a Leica TCS SP5 confocal microscope. Zoom panel shows magnification of IBs boxed in the merge panel. Graphs shows fluorescent intensity profiles along the indicated white lines drawn across one or two IBs. **(C)** MDBK cells were mock infected or infected with bRSV. At the indicated times p.i. cells were fixed and immuno-stained with anti RSV N (green) and anti-NF-κB p65 (red) antibodies. Images are max intensity z-stacks of 8 planes 0.5 μm apart. Cytoplasmic bodies (area >0.1 μm^2^) from the z-stacks were quantified in a total of 18 infected cells per time point as detailed in the methods. **(D)** Number of N and N and p65 positive bodies per cell at the indicated time points. **(E)** Surface area of identified N and N and p65 positive IBs. Statistical significance determined by ANOVA as described in the methods, \*\*\*\**p*<0.0001. **(F)** Vero cells were infected with wt bRSV, ΔNS1, ΔNS2, ΔNS1ΔNS2 or ΔSH bRSV. 24 h p.i., cells were fixed and immunostained with rabbit anti-NF-κB p65 (red) and mouse anti-RSV F (green) antibodies. Cell nuclei were stained with DAPI (blue) and images obtained using a Leica TCS SP5 confocal microscope.

We next examined whether the established bRSV-encoded immunomodulatory proteins - NS1, NS2 [18] and SH [14, 15] - are responsible for this p65 sequestration. We infected cells with wild type bRSV (wt), or recombinant bRSVs which do not express these proteins (ΔNS1, ΔNS2, ΔNS1/2 (a double knockout) or ΔSH - [15, 40]). At 24 h p.i., infected cells were fixed and co-immunostained for p65 and the RSV F protein. To confirm deletion of SH, immunostaining was performed using an anti-SH antibody (supplementary Fig 4A). Since we did not have access to anti-NS antibodies, the genotype of these mutants was confirmed by RT-PCR, on RNA extracted from infected cells targeting the region of NS deletion (supplementary Fig 4B). IF analysis of these samples identified p65 puncta in all infected cells (Fig 3F), suggesting that these bRSV encoded immunoantagonists do not play a significant role in either the formation of IBs or the sequestration of p65 to these structures.

### bRSV IBs are sites of RNA replication but p65 does not specifically co-localise with M2-1 or nascent viral RNA in IB-associated granules (IBAGs)

hRSV inclusion bodies have previously been shown to be the sites of virus transcription and replication [25, 26, 41]. To confirm bRSV IBs are also the site of viral RNA replication, we carried out nascent RNA labelling using 5-ethynyl-uridine (5EU) incorporation. Mock infected MDBK cells, incubated with 5EU for 1 h, revealed, as expected, 5EU incorporation into cellular RNA in the nucleus (Fig 4A; top row). When cellular transcription was inhibited following pre-incubation of mock infected cells with actinomycin D (Act D) for 1 hr this signal was lost. 5EU labelling performed on bRSV infected cells without Act D treatment did not reveal significant evidence for viral replication in IBs; perhaps due to over-representation of cellular RNA synthesis. However, in the presence of Act D, labelled, newly synthesised RNA could only be seen in the N-positive IBs, presumably the result of viral replication. This co-localisation of 5EU incorporation and N-protein within IBs provides strong evidence that bRSV IBs are the sites of viral RNA replication. A more detailed look at the IBs (Fig 4A; inset zoom and line of interest plot - asterisks) revealed partial sub-IB organisation to the RNA found within these structures. Interestingly, a recent study on hRSV IBs identified similar functional compartments within IBs termed inclusion body-associated granules (IBAGs) [26]. These were shown to concentrate newly synthesised viral mRNA and the viral M2-1 protein but not genomic RNA, or the N, P and L proteins. To confirm the presence of IBAGs in bRSV IBs we immuno-stained bRSV-infected cells for M2-1 following nascent viral RNA labelling, observing co-localisation of both these components (Fig 4B). The intra-IB organisation of RNA replication and M2-1 protein into IBAGs appears, therefore, to be a structurally conserved aspect of orthopneumovirus IBs. We next examined the potential co-localisation of p65 with these sites of nascent vRNA localisation (IBAGs). Although we observed partial sub-IB localisation signals for p65, this did not always co-localise with vRNA (Fig 4B) or, in subsequent experiments, M2-1 (Fig 4C). These findings suggest that there are multiple sub-compartments within bRSV IBs, in addition to IBAGs, which potentially carry out a distinct range of functions.

**Figure 4.**
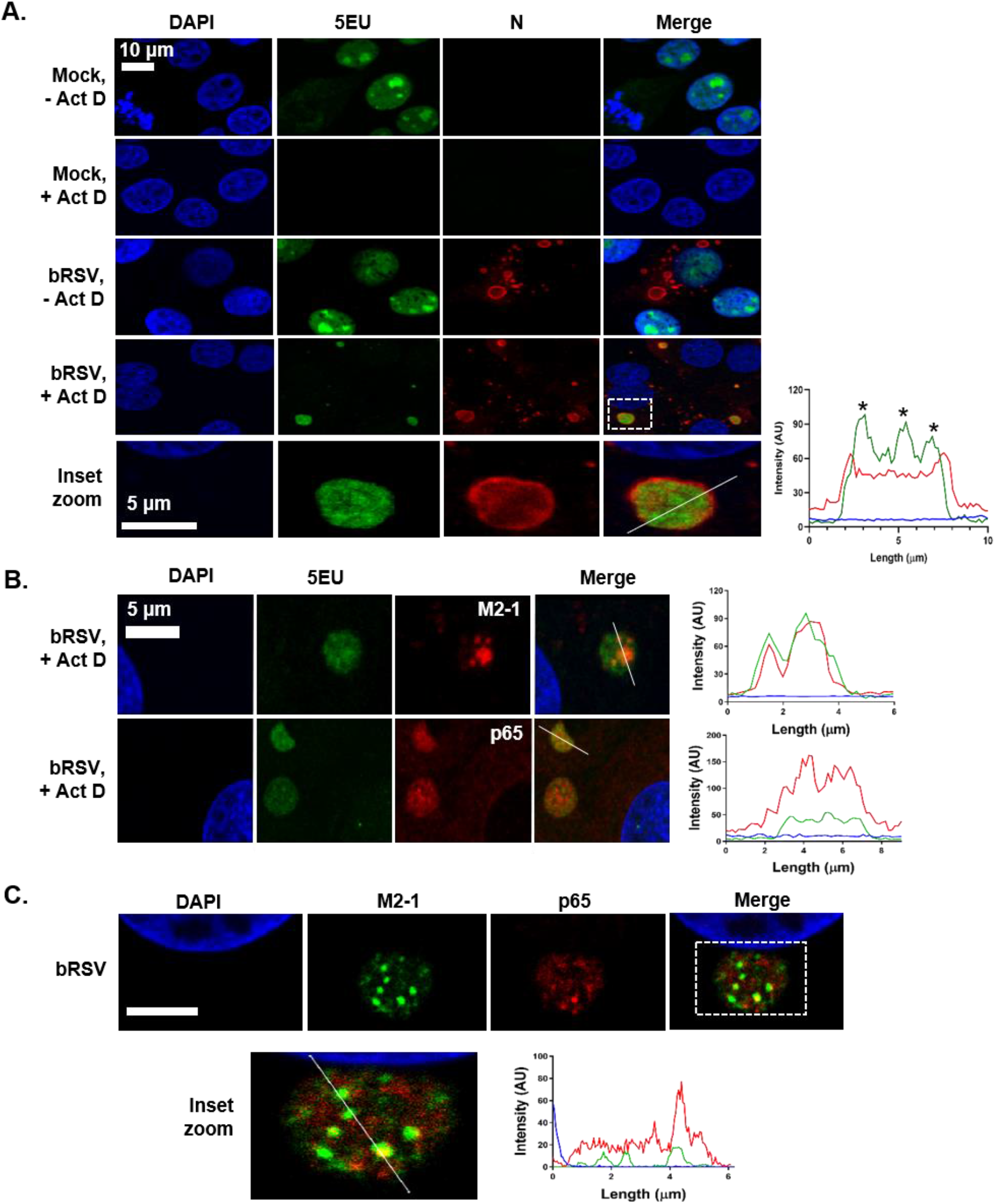
bRSV IBs are sites of RNA replication but p65 does not specifically co-localise with M2-1 or nascent viral RNA in IB-associated granules (IBAGs). **(A)** and **(B)** MDBK cells were mock infected or infected with bRSV. 24 h later, cells were incubated with vehicle or 20 μg/ml actinomycin D (Act D) for 1 h to inhibit cellular transcription. 5-ethynyl uridine (5EU) was then added for another 1 h and the cells fixed. 5EU incorporated into newly synthesised RNA was detected using Alexa Fluor 488-azide (green) as described in the methods. Cells were then immuno-stained with anti-RSV N, M2-1 or anti-NF-κB p65 antibodies (red). Cell nuclei were stained with DAPI (blue) and images obtained using a Leica TCS SP5 confocal microscope. Bottom panel of (A) (inset zoom) shows the boxed area (in merge of bRSV, +Act D) magnified. Graphs show fluorescent intensity profiles along the indicated white lines drawn across the IBs. Asterisks in A. indicate areas of increased 5EU staining within the IB. **(C)** Vero cells infected with bRSV for 24 h were fixed and immuno-stained with rabbit anti-NF-κB p65 (red) and mouse anti-M2-1 (green) antibodies. Cell nuclei were stained with DAPI (blue) and images obtained using a Leica TCS SP5 confocal microscope. Bottom panel shows a higher magnification of the boxed area - scale bar corresponds to 4 μm. Graphs shows fluorescent intensity profiles along the indicated white line.

### bRSV IBs are membraneless liquid organelles

IBs and IB-like structures form by liquid-liquid phase separation (LLPS) which favours macromolecular-macromolecular over macromolecular-water interactions [42–44]. The resulting biomolecular condensates are not surrounded or compartmentalised by a membrane, distinguishing them from many other organelles found in the cytoplasm [44, 45]. To examine the ultrastructural properties of the bRSV IBs we first performed standard transmission electron microscopy (TEM) of infected cells. Vero cells were infected with bRSV at an MOI of 1 and fixed for TEM analysis at 24 and 48 h p.i. Granular structures with high electron density, characteristic of RNA virus inclusion bodies, were identified at both timepoints, often in close proximity to the nucleus (Fig 5A). Smaller structures (1-2 μm in diameter) were predominately rounder in nature when compared to their larger (>3 μm in diameter), more pleomorphic counterparts (Fig 5A). As expected, these structures were not membrane-bound or directly associated with sub-cellular organelles; however, rough endoplasmic reticulum (RER) and mitochondria were frequently found in close proximity (Fig 5A). These structures are similar to those previously reported for other RNA viruses [28, 39], supporting our conclusion that bRSV also forms membraneless IBs in infected cells.

**Figure 5.**
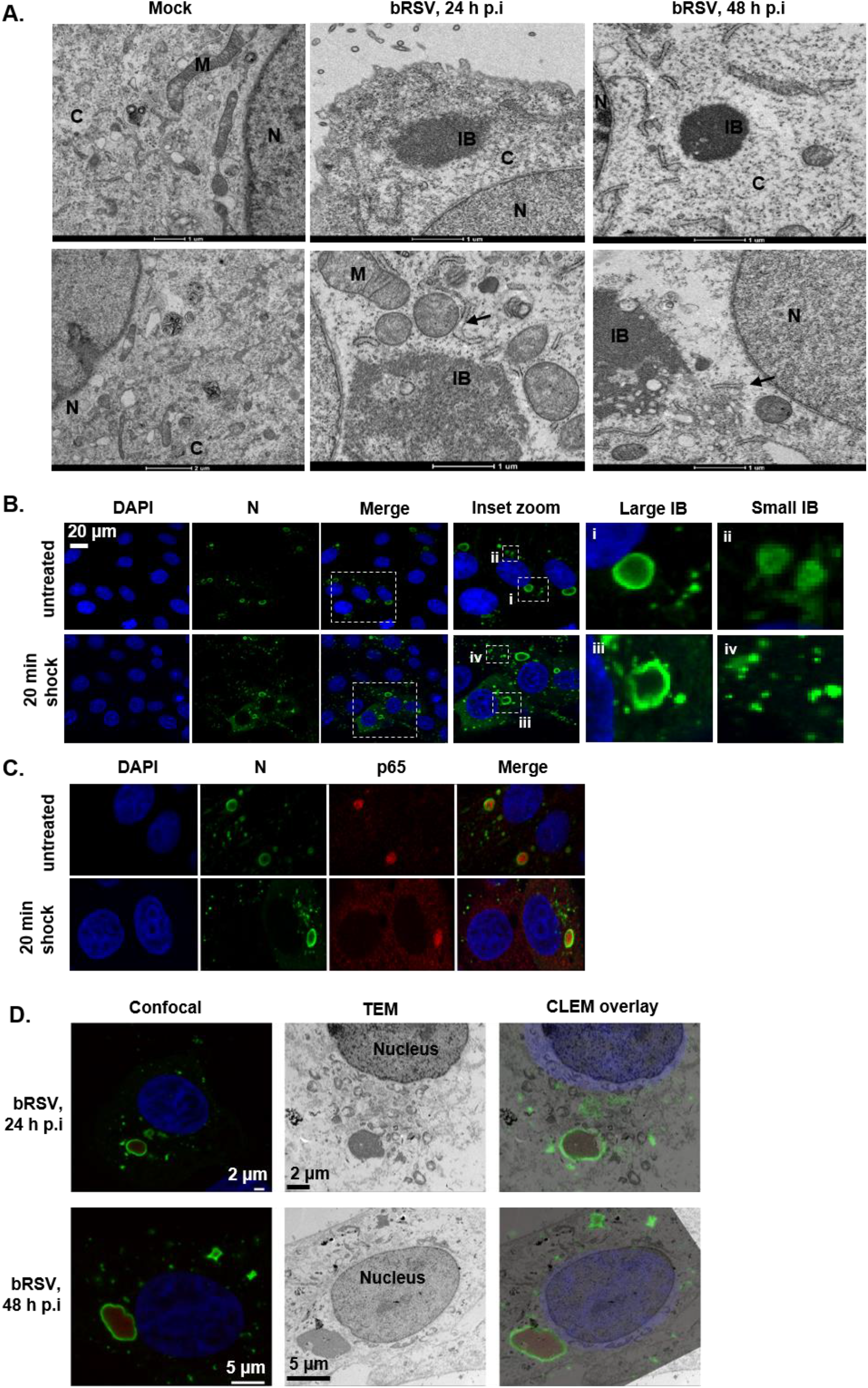
bRSV IBs are membraneless liquid organelles. **(A)** High power transmission electron microscopy (TEM) of mock or bRSV infected Vero cells fixed in glutaraldehyde at 24 and 48 h p.i and prepared for TEM as detailed in the methods. N, nucleus; M, mitochondria; C, cytoplasm; IB, inclusion body and ER indicated with black arrow. Two representative images are shown per time point. Scale bars correspond to 1 μm. **(B/C)** Vero cells were infected with bRSV at an MOI of 1 and incubated at 37°C for 24 h. Hypotonic shock was applied for 20 mins before the cells were fixed. Confocal analysis was performed following immuno-staining for bRSV N (green) and nucleus stained with DAPI (and also p65 for C.). Inset zooms demonstrate the observed effects of hypotonic shock on large (i and iii) and small (ii and iv) IBs – representative images shown. **(D)** Correlative light electron microscopy (CLEM) of confocal microscopy immunostaining and TEM showing bRSV IBs. Vero cells infected with bRSV at MOI 1 were fixed at 24 or 48 h p.i., stained with antibodies against RSV N (green), NF-κB p65 (red) and nuclei stained with DAPI. Following confocal imaging, cells were fixed in glutaraldehyde, sectioned and visualised by TEM. Confocal (left) and TEM (middle) images of the same cells were overlayed (right) as CLEM images.

Various reports have also demonstrated that IBs can rapidly change their size due to fusion or fission whilst remaining spherical in nature, a characteristic feature of these liquid organelles [42]. Rabies virus inclusion bodies, termed negri bodies, have been shown to rapidly dissolve and reform in response to hypotonic shock, demonstrating the dynamic nature of these structures [27, 28, 46]. To assess the sensitivity of bRSV IBs to hypotonic shock, Vero cells, infected with bRSV for 24 h, were incubated with DMEM (diluted to 20% in H_2_O) for 20 mins. Cells were then fixed and immunostained for N protein. Many smaller IBs showed evidence of dissolution following hypotonic shock (Fig 5B; iv); however, unlike rabies virus negri bodies, the larger bRSV IBs remained intact following this significant period of cellular osmotic shock (Fig 5B; iii). Of note, incubation beyond 20 minutes was not possible because of the associated cytotoxicity. In addition, a large percentage of the sequestered p65 in these larger IBs remained tightly associated with the intact structure (Fig 5C). Recently, Zhou et al., demonstrated that larger measles IBs had slower rates of fluorescence recovery after photobleaching (FRAP), relative to their smaller counterparts, postulating that these structures had acquired a more gel-like property. The acquisition of this gel-like status, which are also less likely to exchange molecules with the surrounding cytoplasm, has been linked to aging of phase separated organelles - a continuum which ends with the formation of irreversible aggregates [47]. The insensitivity of large bRSV IBs to osmotic shock, and the maintenance of p65 within the IB even under these harsh conditions, is perhaps the result of them acquiring gel-like status, a property which may be linked to the age and size of individual IBs within infected cells.

Finally, to examine the sub-IB localisation of RSV N and p65 in relation to our ultrastructural analysis of IBs, we performed correlative light electron microscopy (CLEM). Vero cells were infected at an MOI of 1 and analysed at 24 and 48 h p.i., firstly by confocal microscopy using N and p65 antibodies to immunolabel these proteins (Fig 5D). The same cells, identified by grid reference, were then isolated, embedded and sectioned with their ultrastructure subsequently analysed by TEM. Importantly, these CLEM data confirmed that the electron dense granular structures seen by TEM (Fig 5A) are synonymous with the N, P, M and p65 stained IBs seen in IF microscopy (Fig 3B). To our knowledge this is the first CLEM to be performed on an RNA virus IB. An overlay of the two images confirmed that bRSV IBs had retained the electron dense granular structure characteristic of liquid organelles, even with the chemical permeabilization required for IF antibody labelling (Fig 5C and supplementary Fig 5). Our CLEM data also confirmed the p65 and N proteins localising to the IB, with p65 present within the structure and N around the periphery. At 24 h p.i., the p65-positive IB structures were mostly spherical, becoming larger and more irregularly shaped by 48 h p.i., possibly as a result of transition into a more gel-like status, as discussed above. A similar pattern of immunostaining and IB morphology was also observed in bRSV-infected MDBK cells analysed by CLEM (Supplementary Fig 5).

### Co-expression of bRSV N and P proteins induces the formation of IB-like structures which can sequester p65

In the absence of infection, ectopic co-expression of many *Mononegavirales* N and P proteins has been shown to result in the formation of IB-like structures [24, 26, 28, 30] – a finding which has been linked to their potential to induce LLPS independently of viral infection. Although there has been broad discussion that this is related to the presence of intrinsically disordered regions within the N and P proteins, a definitive functional mechanism for this viral-induced LLPS remains uncharacterised. In addition, whether these infection-independent IB-like structures retain all the properties of viral IBs is not entirely clear. For hRSV it was shown that IBAGs do not form within these visually orthologous bodies [26]; however, the recruitment of MDA5 and MAVS to IB-like structures, following N and P overexpression, was maintained [24]. To address similar questions for bRSV IBs, and to examine the related sequestration of p65, Vero cells transiently transfected with plasmids expressing bRSV N (pN) and bRSV P (pP) were fixed and stained at 24 h post transfection and examined by IF. As has been reported previously, expression of N or P alone did not lead to the formation of IB-like structures; however, co-expression did, resulting in the formation of inclusions up to 6.9 μm^2^ in area (Fig 6A). Examination of the sub-cellular localisation of p65 in this system also confirmed that the N- and P-induced inclusions were proficient in sequestering p65, independent of viral replication, with a pattern of expression mirroring that seen in infected cells (Fig 6A; inset zoom and fluorescent line of interest analysis).

**Figure 6.**
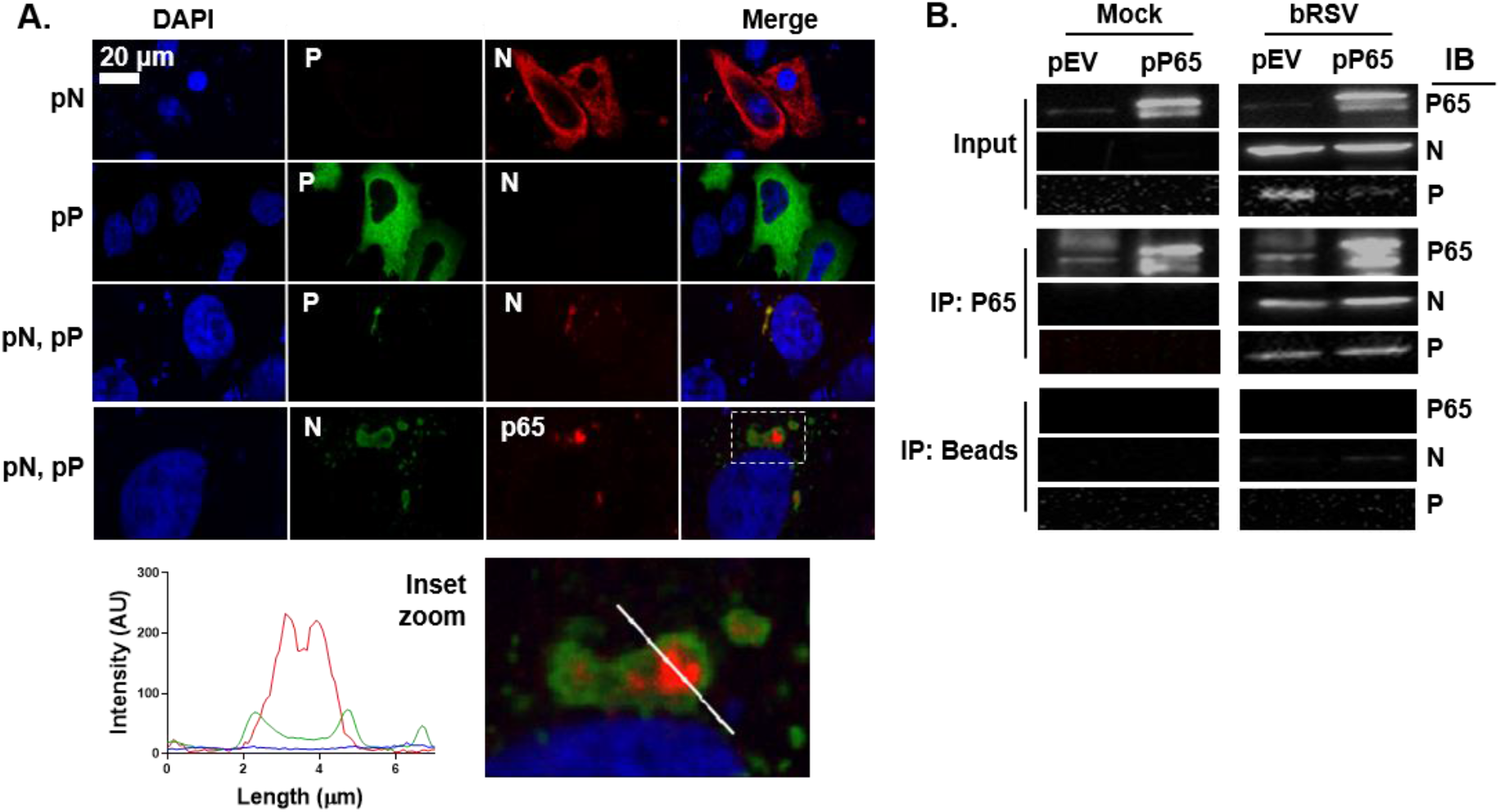
Co-expression of bRSV N and P proteins induces the formation of IB-like structures which can sequester p65. **(A)** Vero cells were co-transfected with equimolar concentrations of plasmids expressing bRSV N (pN) and/or P (pP) proteins as indicated. Following 24 h incubation, cells were fixed and stained with anti-RSV N (green/red) and anti-RSV P (green) or anti-NF-κB p65 (red) antibodies. Bottom panel shows a higher magnification of the boxed area. Graphs shows fluorescent intensity profiles along the indicated white line. **(B)** Co-immunoprecipitation of p65. 293T cells were transfected with plasmids expressing NF-κB p65 (pP65) or empty vector (pEV) and 6 h later infected with bRSV at MOI 1. At 24 h p.i., cell lysates were immunoprecipitated (IP) with anti-p65 antibody or beads alone as a control. Pull-downs were analysed by SDS-PAGE and immuno-blotting (IB) using anti-p65, anti-N or anti-P antibodies

To examine the mechanism of p65 recruitment to, and sequestration within, the bRSV IB we next investigated whether there was evidence for direct protein-protein interactions between this protein and N or P. Endogenous p65 or p65 expressed from a plasmid (pP65) were immunoprecipitated from bRSV-infected, or mock-infected, 293T cells (at 24 h p.i) using an anti-p65 antibody. When these immuno-precipitates were analysed by western blot, both bRSV N and P were found to co-immunoprecipitate (co-IP) with endogenous or overexpressed p65 in infected cell lysates, providing evidence of direct interactions being maintained post-lysis (Fig 6B). Experiments with beads alone did show a small amount of co-IP N protein; however, this was markedly lower than in the p65 antibody experiment, background signal which we believe may be the consequence of the high levels of N protein in infected cells at 24 h p.i. In summary, our results indicate that p65 recruitment into bRSV IBs is maintained even in IB-like structures formed after N and P overexpression. Furthermore, the recruitment of p65 to IBs is likely due to specific interactions with the N and/or P proteins. Since RSV N and P are known to interact, yet the IB does not form without both proteins being expressed together, more detailed characterisation of this interaction is required to define the true binding partner, either N or P.

### The sequestration of the NF-κB subunit p65 to cytoplasmic IBs is a conserved mechanism of orthopneumovirus immunomodulation

Having established structural and functional similarity between bRSV and hRSV IBs, we finally examined the regulation and sub-cellular localisation of the NF-κB subunit p65 in hRSV infected cells. Beginning with the NF-κB luciferase reporter assay we uncovered a pattern of signalling inhibition similar to bRSV. Infection with hRSV in the presence of the NF-κB reporter did not lead to robust activation when compared to mock infected cells, highlighting a lack of activation of this pathway in infected cells (Fig 7A, black bars). Again, similar to bRSV, infected 293T cells (24 h with hRSV) which were stimulated for 6h with hTNFα induced significantly less NF-κB transactivation, when compared to equivalently treated mock-infected cells (Fig 7A, grey bars). This correlated well with an examination, by IF, of hRSV replication in Vero cells, with and without hTNFα treatment, where again we did not observe significant levels of p65 nuclear translocation (Fig 7B). Indeed, as observed in bRSV infected cells, p65 was recruited into intra-cytoplasmic puncta. These puncta were subsequently shown to be synonymous with viral IBs (Fig 7C) in a set of experiments which also confirmed that IB formation and the recruitment of p65 is host cell independent. bRSV or hRSV infected MDBK (bovine) or Hep2 (human) cells demonstrated the presence of p65-containing IBs in all scenarios, highlighting that the mechanisms underpinning RSV IB formation, and the sequestration of p65 to these bodies, are likely highly conserved (Fig 7C). We concluded this examination of host-range specificity with a more physiologically relevant model of the human bronchial epithelium, BEAS-2B cells, which are derived from normal human tissues taken following autopsy of non-cancerous individuals, identifying again the formation of IBs and sequestration of p65, regardless of RSV species. Finally, we confirmed that IB-like structures formed by ectopic hRSV N and P co-expression recruited p65 to their core (Fig 7D). Taken together, these data indicate that the formation of IBs during viral replication, together with the sequestration of the transcription factor NF-κB subunit p65 to these bodies, is a common feature of orthopneumoviruses.

**Figure 7.**
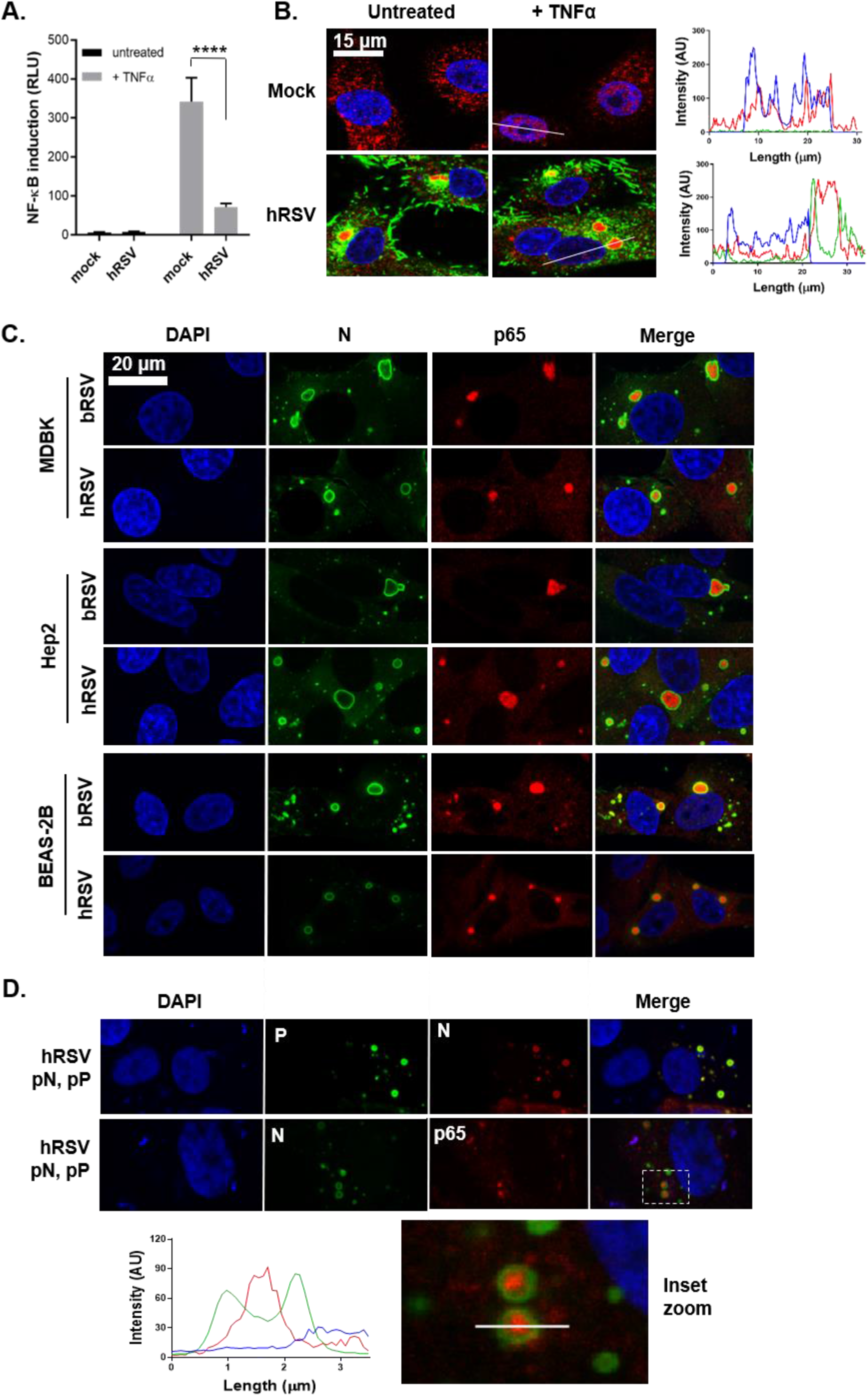
The sequestration of the NF-κB subunit p65 to cytoplasmic IBs is a conserved mechanism of orthopneumovirus immunomodulation. **(A)** 293T cells were mock infected or infected with hRSV at an MOI of 1. At 6 h p.i., cells were transfected with 100 ng NF-κB FLuc reporter and 10 ng TK-renilla luciferase and incubated at 37°C. At 18 h p.t., cells were left untreated or stimulated for 16 h with 20 ng/ml hTNFα. Cells were then lysed and analysed for firefly and renilla luciferase activities. Graph depicts means ± SD of three replicates from the same experiment. Statistical significance determined by ANOVA as described in the methods, \*\*\*\**p*<0.0001. **(B)** Vero cells mock infected or infected with hRSV at an MOI of 1 for 24 h were left untreated or stimulated with 20 ng/ml hTNFα for 30 mins. Cells were then fixed and immuno-stained with anti-RSV F (green) or anti-NF-κB p65 (red) antibodies. Cell nuclei were stained with DAPI (blue) and images obtained using a Leica TCS SP5 confocal microscope. Graphs show line fluorescent intensity profile along the indicated white lines. **(C)** MDBK, Hep2 and BEAS-2B cells were infected with b/hRSV for 24 hrs, fixed and immuno-stained for RSV N (green) or NF-κB p65 (red). **(D)** Vero cells were co-transfected with equimolar concentrations of plasmids expressing hRSV N (pN) and/or P (pP) proteins as indicated. Following 24 h incubation, cells were fixed and stained with anti-RSV N (green/red) and anti-RSV P (green) or anti-NF-κB p65 (red) antibodies. Cell nuclei were stained with DAPI (blue) and confocal analysis performed. The bottom panels show a higher magnification of the boxed area and a graph with the fluorescent intensity profiles along the indicated white line.

## Discussion

Recognition of viral pathogen-associated molecular patterns (PAMPs) by RIG-I or MDA-5 can lead to activation of NF-κB transcription factors through the IKK complex or IRFs through TBK-1/IKKε [9, 11, 48]. Activation of these innate responses is essential for inducing a robust adaptive response, firstly to clear viral infections and secondly to elicit the establishment of a memory response [4, 48]. However, in vivo the various immune-evasion strategies employed by RSV combine to generate only a short-lived response [4, 19, 20, 23, 48]. For instance, there is strong evidence that the downregulation of key signalling molecules by the NS proteins suppresses IRF3 activation and type I IFN induction [17–20, 22, 23], although interestingly we did see significant IRF3 nuclear translocation in our infected cells. As a key innate immune pathway, NF-κB signalling is often a target for viral antagonism; however, to date RSV modulation of its activation has remained less well defined. Although RSV lacking the *SH* gene was shown to enhance NF-κB activation, the exact mechanisms employed are unclear [14, 15, 49, 50]. To address this, we monitored NF-κB p65 activation in RSV-infected cells at multiple steps in the signalling pathway: IκBα degradation, p65 phosphorylation (at Ser536), p65 nuclear translocation, and more broadly NF-κB transactivation. We present a novel mechanism of immune evasion wherein RSV infection results in the sequestration of the NF-κB subunit p65 into viral inclusion bodies (Fig 3B), a process which is independent of the known RSV immunomodulatory proteins, NS1, NS2 and SH (Fig 3F). We also demonstrate that as a result, activation of NF-κB p65 is suppressed in infected cells, even with exogenous TNFα stimulation (Fig 1). Although small IBs were observed as early as 6 h p.i. (≤2.5 μm^2^) these did not colocalise with detectable levels of p65 (Fig 3E). This may reflect a technical limitation of our IF, or alternatively that IBs need to grow in size before they can begin to sequester p65. It remains to be determined if p65 is actively recruited to IBs by viral proteins or if its sequestration is a result of the IB’s position in the cell and that it captures p65 by an indirect mechanism, perhaps involving trafficking. Interestingly, the lack of p65 activation prior to IB formation and p65 aggregation, highlights that RSV may employ additional mechanisms for NF-κB inhibition which remain uncharacterised. From a wider perspective, this mechanism of immunomodulation might be a common strategy utilised by RSV and other viruses that induce IB formation. MAVS and MDA5 were similarly both found to be recruited into RSV IBs as a mechanism of suppressing IFN signalling [24]. Similarly, p38 MAPK and OGT sequestration into RSV IBs suppressed MAPK-activated protein kinase 2 signalling and stress granule formation, respectively, enhancing virus replication [25]. Whether viruses such as Ebola, Nipah or rabies adopt similar mechanisms of immunomodulation remains to be determined.

From a mechanistic perspective our results also showed that the N and P proteins are essential for the formation of bRSV IBs. As has been reported for rabies [28] and measles [30] viruses, ectopic expression of these proteins resulted in the formation of IB-like structures (Fig 6A and 7D). These were mostly spherical and at 24 h post transfection, measured up to 6.9 μm^2^ which is considerably less than the conventional IBs observed in infected cells. We hypothesise that both pseudo-IBs and viral IBs form by biomolecular condensation, but that their maturation into larger structures is dependent on other factors present only in infected cells. That these pseudo-IBs could also recruit p65 suggested a direct interaction between p65 and RSV N or P, which we confirmed by co-IP (Fig 6B). Interestingly, our IF data was somewhat contradictory, with the staining patterns and line intensity profiles showing p65 concentrated in the middle of IBs with N and P at the periphery, separating the IB contents from the cytoplasm. It is possible that exchange of biomolecules across the boundary, e.g. during the sequestration of p65, may require transient N or P interactions. Intriguingly, Lifland et al. also suggested MAVS and MDA-5 are recruited into IBs by interacting with N and P in a macromolecular complex [24]. We propose that this recruitment may involve low-affinity interactions with N and/or P and that maintenance within the IB is enhanced by the same physicochemical properties of the IBs which enable them to induce LLPS, namely macromolecular-macromolecular interactions. The RSV P protein has been shown to bind and recruit M2-1 to IBs, potentially through intrinsically disordered regions within P that allow it to form multiple interactions [51]. Although further work is required to identify the exact mechanism of p65, MAVS, MDA5 etc. recruitment into IBs, we postulate the physicochemical properties of these proteins may also be an important factor.

Electron micrograph analysis of our RSV IBs showed greater electron density in the IBs, when compared to the cytoplasm, a characteristic of biomolecular condensates (Fig 5A). These data also highlighted the structural complexity of the phase-separated structure. Although we observed some association with the ER and RER, RSV IBs were not membrane bound, unlike rabies virus negri-bodies which acquire a membrane boundary later in infection, presumably derived from the ER [28, 39]. Interestingly, our CLEM analysis confirmed previous IF data from the field that the IB boundary is surrounded by N protein (Fig 5D). A debate remains in the field as to whether this is an artefact of disrupted antibody epitope accessibility to N, since GFP tagged N proteins were shown to have a diffuse pattern throughout the IB [24]; however, we would only note that we used an antibody developed in-house for this staining. Nevertheless, the presence of viral RNA associated proteins, N, P and M2-1, in IBs (Fig 3B and 4C) strongly suggested the presence of RNA replication and transcription within these structures. Building on previous work for hRSV and rabies virus [26, 39], we used 5EU incorporation to confirm RNA synthesis in the IBs (Fig 4A and B). Using fluorescence in situ hybridization (FISH) experiments, Rincheval et al. showed that genomic RNA colocalised with the hRSV N and P proteins at the periphery, whilst viral mRNA was found to concentrate in IBAGs, transient sites of mRNA storage [26]. Our data showed the formation of similar structures, confirming IBAGs are found in multiple orthopneumoviruses; however, there was no conclusive colocalisation with p65. However, this sequestered cellular protein did localise to distinct intra-IB bodies (Fig 4B and C), raising the intriguing possibility that multiple microdomains exist within what is, by TEM, an apparently uniform granular biomolecular condensate.

In summary our data shows that RSV IBs are highly ordered structures performing multiple roles in the virus lifecycle including the compartmentalisation of virus replication and transcription and the sequestration of cellular proteins involved in the antiviral response. This mechanistic characterisation is potentially applicable to other negative sense RNA viruses that have been shown to form IBs during replication.

## Materials and Methods

### Cells and viruses

All cells were cultured at 37ΰC in a 5% CO_2_ atmosphere. MDBK (Madin-Darby bovine kidney), Vero (monkey kidney epithelial), 293T (human embryonic kidney) and Hep-2 (human epithelial type 2) cells were obtained from the Pirbright Institute Central Services Unit and maintained in Dulbecco’s Modified Eagle’s Medium (DMEM) supplemented with 10% heat inactivated foetal calf serum (FCS; TCS Biologicals), sodium pyruvate (Gibco), penicillin and streptomycin (Sigma). Beas-2B (human bronchial epithelial) cells (ATCC) were cultured in LHC basal medium (ThermoFisher) supplemented with 10% FCS, penicillin and streptomycin.

Wild-type recombinant (r) bRSV and deletion mutant rbRSVs ΔSH, ΔNS1, ΔNS2, and ΔNS1/2 were produced by reverse genetics from rbRSV strain A51908 variant Atue51908 (GenBank accession no. AF092942) [18, 40, 52]. These were propagated in Vero cells and hRSV subtype A (A2 strain) grown in Hep-2 cells. All viruses were further purified from total cell lysates using polyethylene glycol (molecular weight, 8,000) precipitation and discontinuous sucrose gradient centrifugation.

### Plasmids and transfections

All viral gene sequences were derived from bRSV A51908 (GenBank accession NC_038272) and hRSV A2 (GenBank accession KT992094). Expression plasmids (pcDNA3.1) encoding codon-optimised *N* genes at KpnI-BamHI sites referred to as pN were purchased from Bio Basic Inc. Full length *P* genes were amplified by reverse transcriptase PCR using gene-specific primers and Superscript II reverse transcriptase (Invitrogen). These were then cloned into pcDNA3.1 at KpnI-BamHI sites and designated pP. The p65 open reading frame (ORF) was amplified from pcDNA3.1-HA-p65 (kindly provided by Carlos Maluquer de Motes, Uni. Of Surrey) and inserted at the HindIII-BamHI sites of pcDNA3.1; designated pP65. All sequences were confirmed by conventional sanger sequencing. Plasmids were transfected into cells using TransIT-X2 (Geneflow).

### Antibodies and drugs

Mouse monoclonal antibodies raised against bRSV F (mAb19), N (mAb89), P (mAb12), M (mAb105) and M2-1 (mAb91) were previously described [53, 54]. Rabbit polyclonal anti-bRSV SH antibody was purchased from Ingenasa. Rabbit anti-NF-kB p65 (8242) antibody; rabbit anti-IRF3 (11904); rabbit anti-phospho-NF-kB p65 (Ser536; 3033); mouse anti-κBα (4814); and rabbit anti-GAPDH (5174) were obtained from Cell Signaling Technology. Mouse anti-G3BP-1 was obtained from BD Biosciences. Secondary horse-radish peroxidase-linked antibodies were obtained from CST and Alexa Fluor secondary antibodies from Life Technologies. Recombinant hTNFα (CST), poly(I:C) (InvivoGen), sodium arsenite (Sigma), and actinomycin D (Sigma) were purchased from the indicated suppliers.

### Confocal immunofluorescence microscopy

Cells were fixed with 4% paraformaldehyde (PFA; Sigma) in PBS for 15 mins, permeabilised with 0.2% Triton X-100 in PBS for 5 mins and blocked with 0.5% bovine serum albumin (BSA) (Sigma) in PBS. Cells were then incubated with the indicated primary antibodies overnight at 4°C. They were then washed and incubated with Alexa Fluor secondary antibodies (Life Technologies) for 1 hr at room temperature. Cells were then washed and mounted with Vectashield (Vector labs) containing DAPI for nuclei staining. Fluorescence was imaged on a Leica TCS SP5 confocal microscope using 405nm, 488nm and 568nm laser lines for the appropriate dyes and a 63X oil immersion objective.

### Quantitation of bRSV induced p65 puncta and IBs

Mock or bRSV-infected (at an MOI of 1) MDBK cells were fixed in 4% PFA (Sigma) at 6, 16 and 24 h p.i., and labelled according to the described immunofluorescence method. Multiple Z-sections, 0.5 μm apart, were taken for each cell, by confocal microscopy and max intensity Z-stacks of 8 planes made using the Leica LAS AF Lite software. Quantifications of N and p65 positive structures were performed using the area region of interest analysis tool. GraphPad Prism 7 was used to perform parametric one-way analysis of variance (ANOVA) and Tukey’s multiple comparison tests. ImageJ was also used to make 3D projections of 9 images 0.9 μm apart.

### Luciferase reporter assay

2 x 10^5^ 293T cells seed into 24 well plates a day prior were mock infected, infected with bRSV or hRSV at an MOI of 1. 6 h later, cells were co-transfected with 100 ng NF-κB FLuc reporter which expresses the firefly luciferase gene under the control of five NF-κB repeated transcription factor binding sites and 10 ng TK-ren control plasmid (both kindly provided by Gareth Brady; The Uni. Of Dublin) using Transit-X2 (Geneflow). 24 hours later, cells were stimulated with 20 ng/mL hTNFα for 16 hours or were left untreated. Cells were then lysed with reporter lysis buffer (Promega) and lysates used to determine firefly and renilla luciferase activities on a Glomax luminometer using luciferase assay system (Promega) and coeleterazine (Promega), respectively. Firefly data was normalised to renilla which was used as an internal control of transfection. GraphPad Prism 7 was used to perform parametric one-way analysis of variance (ANOVA) and Tukey’s multiple comparison tests.

### Western blot analysis

Following virus infection and stimulation, growth media was removed from cells and cell extracts prepared by lysis in SDS sample buffer (Bio-Rad) supplemented with β-mercaptoethanol (Sigma), complete mini-EDTA-free protease inhibitors (Roche) and 1 mM sodium orthovanadate (New England Bio-Labs). Lysates were then boiled for 5 mins and 30 μl resolved by SDS PAGE on a 12% polyacrylamide gel and proteins transferred to polyvinylidene difluoride (PVDF) membranes (ThermoScientific). After blocking for 1 hr with 5% dry semi-skimmed milk in 0.1% PBS Tween 20 (PBS-T) membranes were washed with PBS-T and incubated with primary antibodies overnight at 4°C. After washing, the membranes were incubated with the corresponding horseradish peroxidase-conjugated secondary antibodies (CST). Protein bands were detected using Clarity Western ECL substrate (Bio-Rad) and imaged with Bio-Rad ChemiDoc™ MP Imaging System.

### 5-ethynyl uridine (5EU) labelling

Infected cells growing on coverslips were incubated with or without medium supplemented with 20 μg/ml actinomycin D (Act D) to inhibit cellular transcription for 1 h. Cells were then incubated with medium containing 1 mM 5EU and 20 μg/ml Act D for another hour. Medium was then washed off and cells fixed in 4% PFA for 15 mins. Cells were then washed with PBS and permeabilized with 0.2% Triton X-100 for 5 mins. These were both supplemented with 0.125 U/ml RNase inhibitor (Promega). Incorporated 5EU was labelled using the Click-IT RNA Imaging Kit (Invitrogen) following the manufacturer’s protocol. Following that, immunofluorescence staining was done as described above.

### TEM

Cells seeded onto Thermanox coverslips (Thermo Scientific) were fixed at 24 h and 48 h p.i in phosphate buffered 2% glutaraldehyde (Agar Scientific) for 1 hour followed by 1 hour in aqueous 1% osmium tetroxide (Agar Scientific). Following dehydration in an ethanol series; 70% for 30 min, 90% for 15 min and 100% three times for 10 min, a transitional step of 10 min in propylene oxide (Agar Scientific) was undertaken before infiltration with 50:50 mix of propylene oxide and epoxy resin (Agar Scientific) for 1 hour. After a final infiltration of 100% epoxy resin for 1 hour, the samples were embedded and polymerised overnight at 60°C. 80μm thin sections were cut, collected onto copper grids (Agar Scientific) and grid stained using Leica EM AC20 before being imaged at 100kV in a FEI Tecnai 12 TEM with a TVIPS F214 digital camera.

### CLEM

Cells seeded onto gridded glass coverslips (MatTek) were fixed at 24 h and 48 h p.i in 4% PFA (Sigma) and labelled according to the described immunofluorescence method. Selected grid squares were imaged on a Leica TCS SP8 confocal using 405nm, 488nm and 568nm laser lines for the appropriate dyes. The cells were then fixed in phosphate buffered 2% glutaraldehyde (Agar Scientific) for 1 hour followed by 1 hour in aqueous 1% osmium tetroxide (Agar Scientific). Following 15min in 3% uranyl acetate (Agar Scientific), the cells were dehydrated in an ethanol series; 70% for 30 min, 90% for 15 min and 100% three times for 10 min. After infiltration of 100% epoxy resin for 2 hours, the samples were embedded and polymerised overnight at 60°C. The glass coverslips were removed with liquid nitrogen and the appropriate grid squares located. 80μm thin sections were cut, collected onto copper grids (Agar Scientific) and grid stained using Leica EM AC20. The specific cells imaged in the confocal were identified and imaged at 100kV in a FEI Tecnai 12 TEM with a TVIPS F214 digital camera.

### Co-Immunoprecipitation

1 ×10^5^ 293T cells cultured overnight in 12-well plates were transfected with pcDNA3.1-empty vector (pEV) or pcDNA3.1-p65 (pP65) using TransIT-X2 (Geneflow). 24 h later, cells were infected with bRSV at MOI 1 or mock infected and incubated for another 24 h. Cells were then lysed on ice with RIPA lysis buffer (EMB Millipore) and cell debris removed by centrifugation. Cell lysates precleared with protein A coated magnetic beads (CST) were incubated with rabbit antip65 antibodies overnight at 4°C. Lysates were then incubated with protein A coated magnetic beads for 20 mins at room temperature with rotation. Following five washes with PBS-T, immunoprecipitates were eluted with Laemmli sample buffer and subjected to SDS-PAGE and western blot analysis as already described.

## Ethics statement

This research did not use any primary human or animal tissue. BEAS-2B cells were procured from ATCC.

## Author contributions

FJ performed all experiments, apart from the EM and CLEM which were performed by JS and PH. EG was involved in training FJ. FJ and DB analysed the data, designed the experiments, compiled the figures and wrote the manuscript.

## Acknowledgements

This work was supported by a UK Research and Innovation (UKRI; www.ukri.org) Medical Research Council (MRC) New Investigator Research Grant to DB (MR/P021735/1) as well as a UKRI Biotechnology and Biological Sciences Research Council (BBSRC, www.ukri.org) Institute Strategic Programme Grant (ISPG) to The Pirbright Institute and DB (BBS/E/I/00007034 and BBS/E/I/00007030). The funders had no role in study design, data collection and analysis, decision to publish, or preparation of the manuscript. We would like to acknowledge the support of Geraldine Taylor (The Pirbright Institute), Ursula Buchholz (NIAID, NIH), Karl-Klaus Conzelmann (Max-von-Pettenkofer Institut), Andrew Broadbent (The Pirbright Institute), Gareth Brady (Trinity College, Dublin), Carlos Maluquer de Motes (University of Surrey) and Helena Maier (The Pirbright Institute) for the provision of valuable reagents, recombinant viruses and technical advice.

