## Supplemental figures for "Respiratory syncytial virus sequesters NF-κB subunit p65 to cytoplasmic inclusion bodies to inhibit innate immune signalling"

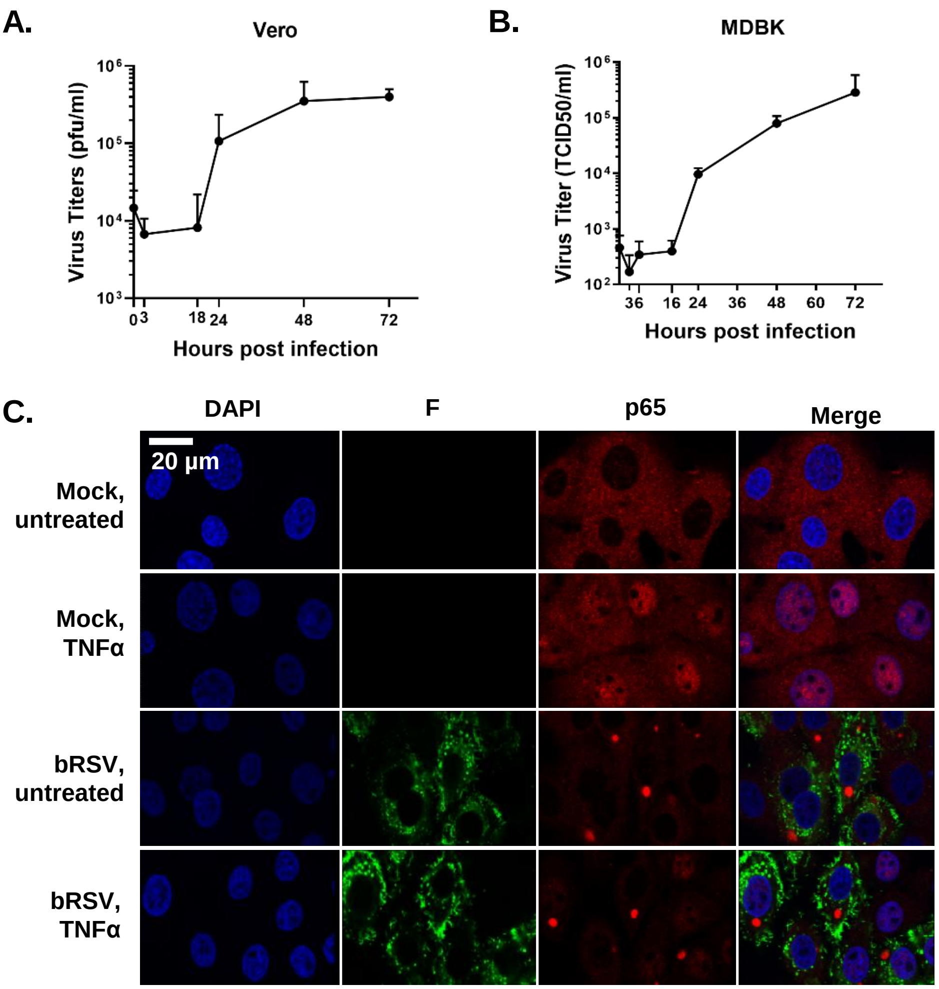


**S1 Fig. bRSV growth and effects on NF-κB activation.** For virus replication, Vero **(A)** and MDBK **(B)** cells were infected with bRSV at an MOI of 2. Viruses were harvested by snap freezing at the indicated times p.i, and virus titres determined by TCID50 end point assay. Graph shows means ± SD of triplicate infections from the same experiment. **(C)** NF-κB activation in bRSV infected MDBK cells. MDBK cells mock infected or infected with bRSV at an MOI of 1 for 24 h were left untreated or stimulated with 20 ng/ml hTNFα for 30 mins. Cells were then fixed and immuno-stained with anti-RSV F (green) or anti-NF-κB p65 (red) antibodies. Cell nuclei were stained with DAPI (blue) and images obtained using a Leica TCS SP5 confocal microscope.

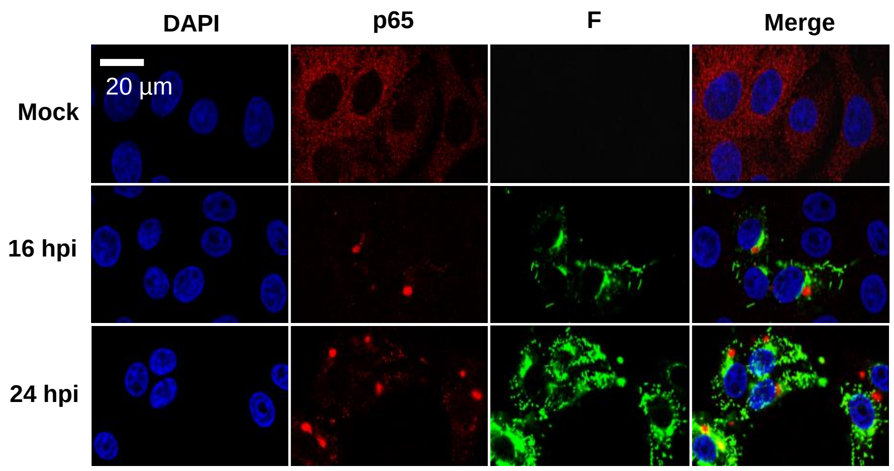


**S2 Fig. Time course of p65 puncta formation in bRSV infected Vero cells.** Vero cells were mock infected or infected with bRSV. At the indicated times p.i., cells were fixed and immuno-stained with anti-NF-κB p65 (red) and anti-RSV F (green) antibodies. Nuclei were stained with DAPI (blue) and images obtained using a Leica TCS SP5 confocal microscope.

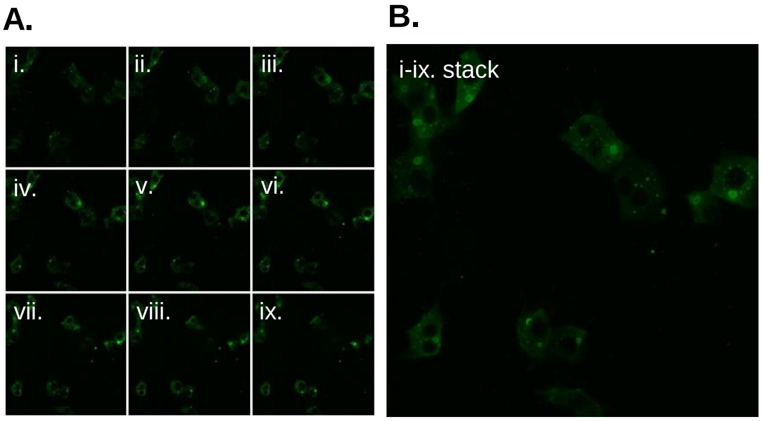


**S3 Fig: Z-stack imaging of RSV IBs.** MDBK cells infected with bRSV were fixed and immunostained with anti-RSV N. Mutiple Z-section images, 0.9 µm apart, were taken through the cell to allow quantification of N positive puncta. In the above example, 9 images (**A**), from cells infected for 24 h, were used to generate Z-stacks (**B**) and Z-projections to quantify N positive IBs and to render 3D projections of these cells (please see supplementary movie files).

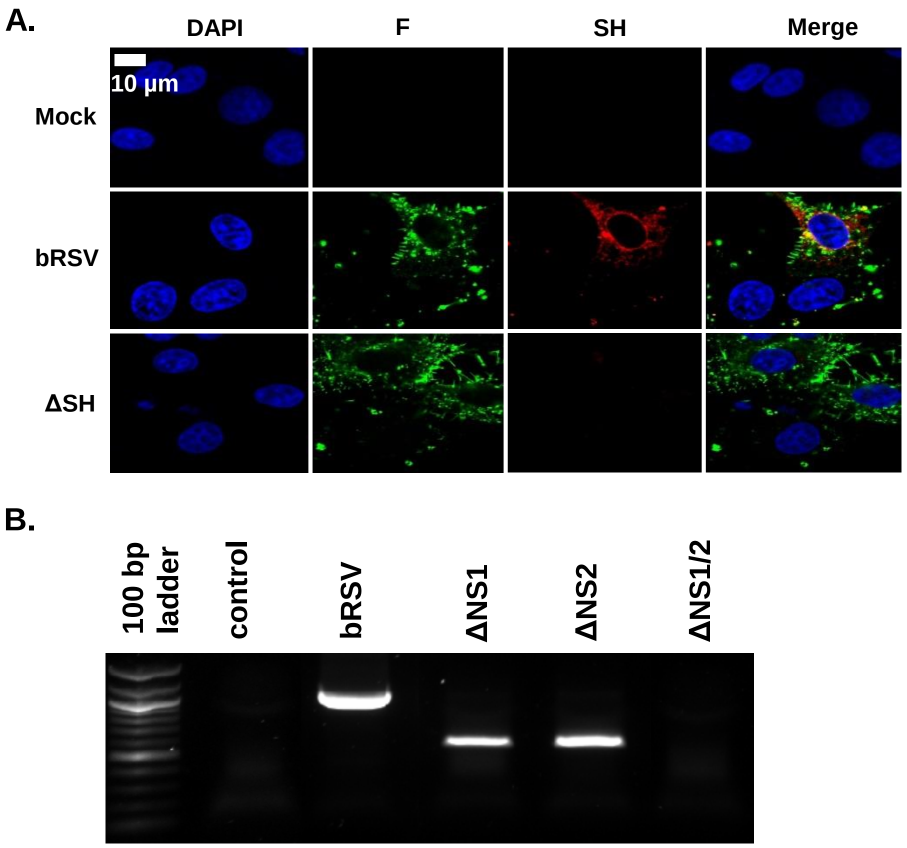


**S4 Fig. Confirmation of bRSV NS1, NS2 and SH gene deletions from recombinant viruses. (A)** Vero cells were mock infected, infected with bRSV wt or ΔSH at an MOI of 1. At 24 h p.i., cells were fixed and immuno-stained with anti-RSV F (green) or polyclonal anti-SH antibodies. Nuclei were stained with DAPI (blue) and images obtained using a Leica TCS SP5 confocal microscope. **(B)** Total RNA was extracted from Vero cells infected with the indicated viruses at 24 h p.i. and subjected to reverse transcriptase PCR using primers that amplify from the gene start (Gs) region of *NS1* to the beginning of the *N* gene.

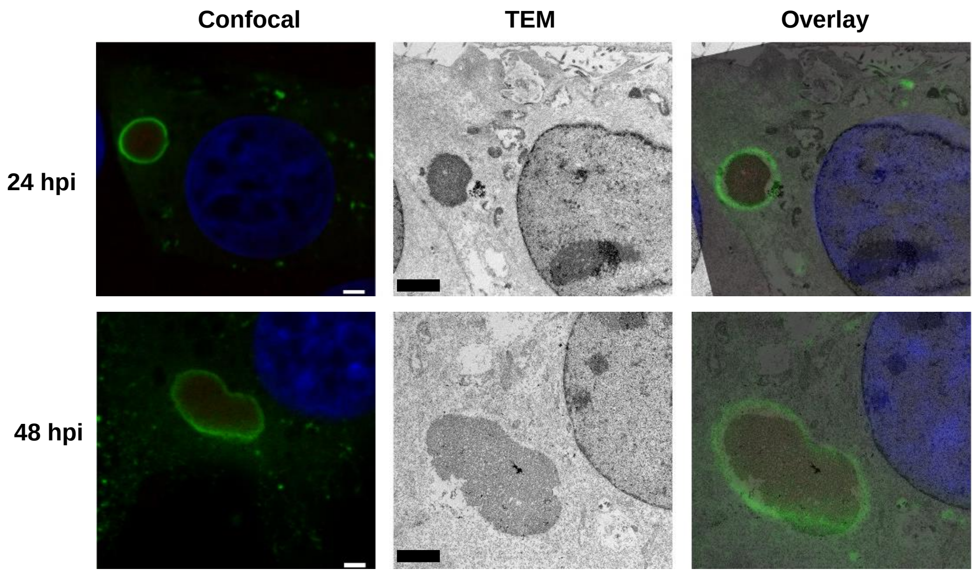


**S5 Fig. Ultrastructural analysis of bRSV infected MDBK cells by CLEM.** MDBK cells infected with bRSV at MOI 1 were fixed at 24 or 48 h p.i., stained with antibodies against RSV N (green), NF-κB p65 (red) and nuclei stained with DAPI. Following confocal imaging cells were fixed in glutaraldehyde, sectioned and visualised by TEM. Confocal (left) and TEM (middle) images of the same cells are overlayed (right) as CLEM images. Scale bars corresponds to 2 µm.
